# Comparison of the NeuMoDX, Diasorin Simplexa, Cepheid and Roche CDC SARS-CoV 2 EUA assays using nasopharyngeal/nasal swabs in universal transport media (UTM) and sputum and tracheal aspirates

**DOI:** 10.1101/2020.05.26.118190

**Authors:** Robert Tibbetts, Kathy Callahan, Kareem Rofoo, Richard J. Zarbo, Linoj Samuel

## Abstract

In March 2019 the outbreak of SARS-CoV 2 was officially defined as a pandemic by the World Health Organization and shortly after, the United States Food and Drug Administration (FDA) granted Emergency Use Authorization (EUA) to the Centers for Disease Control (CDC) for reverse transcription polymerase chain reaction (rtPCR) molecular testing for the detection of the SARS-CoV-2 virus from NP swabs. Since then, EUA with relaxed regulations were granted to numerous manufacturers and clinical microbiology laboratories to implement in-house testing assays with nasopharyngeal swabs (NP) and subsequently additional specimen types. Because of supply chain shortages leading to competition for reagents, sustaining any significant volume of testing soon became problematic. As a countermeasure, within several weeks the Henry Ford Microbiology Laboratory validated 4 different rtPCR assays and multiple specimen types using NeuMoDX, Diasorin Simplexa, Cepheid and Roche platforms. The purpose of this study was to analyze the analytic sensitivity of these rtPCR assays with NP/nasal swabs and sputum/tracheal aspirates. Qualitative analytic agreement between the 4 platforms for NP/nasal swabs ranged 95% - 100% overall with no statistically significant difference in threshold cT values. Similar results were obtained with the sputum/tracheal aspirates. These data demonstrate the high accuracy and reproducibility in detection of SARS-CoV 2 between the rtPCR assays performed on 4 different platforms with numerous specimen types.

## Introduction

As of May 22^nd^, the COVID-19 pandemic that originated in Wuhan, China in December 2019 has infected over 5 million people world-wide resulting in over 335,000 deaths. Of these, 1.6 million total cases and 96,526 deaths have been reported within the United States (https://www.worldometers.info/coronavirus/#countries, accessed 5/22/2020).

SARS-CoV-2 is a member of the betacoronavirus clade of the Coronaviridae family and is more closely related to SARS-CoV-1 and MERS, which share a common zoonotic origin (1). However, SARS-CoV-2 does share some similarities to the alphacoronavirus clade, which typically cause up to 30% of common colds and some gastrointestinal diseases (1). Coronaviruses are small (120 nm diameter), spherical virons surrounded by the spike or “S” receptor binding proteins, resulting in a “corona” appearance in electron microscopic images. Genomically, these virons consist of 27-34 kb of single-stranded, positive-sense RNA encoding ~ 16 non-structural proteins and 5 structural proteins: spike (S), nucleocapsid (N), a phosphoprotein (M), hemagglutinin/esterase (HE), and a non-glycosylated membrane protein (E) (1). The HE protein is only found in the betacoronavirus clade and is functionally analogous to the HE gene found in Influenza C virus (1).

SARS-CoV-2 viral loads have been demonstrated to peak within 1 week of symptom onset and that the spectrum of shedding can vary considerably based on the severity of illness. Therefore, it is crucial for diagnostic assays to use highly sensitive molecular methods to detect SARS-CoV-2 across a broad spectrum of viral loads (2). Within several weeks of outbreak in the United States, the Food and Drug Administration (FDA) gave Emergency Use Authorization (EUA) to the CDC to develop a PCR based assay to detect the SARS-CoV-2 specific N1 and N2 gene sequences (https://www.fda.gov/news-events/press-announcements/coronavirus-covid-19-update-fda-issues-new-policy-help-expedite-availability-diagnostics). Subsequently, numerous manufacturers have also received EUA status for in vitro diagnostic devices (IVD) (3). With national demand outstripping supply, reagent supplies for all available test platforms as well as suitable nasopharyngeal (NP) and nasal collection swab devices became severely limited early on and forced clinical laboratories to quickly validate additional options for specimen collection and testing. The purpose of this study was to compare the performance of three rtPCR testing platforms with a panel of positive and negative samples using NP/nasal swabs and sputum/tracheal aspirates in a modified CDC-based assay for SARS-CoV-2 on Roche LC480, Diasorin Simplexa (DIA), NeuMoDX (NDX) and Cepheid GenXpert (GX). The NDX is a relatively novel, high throughput system that was undergoing preclinical testing prior to the pandemic and has no FDA approved IVD assays to date whereas the DIA and GX systems are well established and have a large menu of IVD assays. Each of these platforms offer different options in terms of workflow, throughput and sample handling that will also be discussed.

## Materials and Methods

### Patient specimens

NP/Nasal swabs in universal transport media (UTM): A total of 20 positive and 30 negative NP/UTM patient specimens tested using the modified CDC PCR were used in this study. Specimens were processed for each assay according to manufacturer’s direction as described below. Ten of the positive specimens were tested on the additional assays within 24 - 48 hours of initial testing and the remaining 10 positives were tested within 72 hours of initial testing with the CDC assay. All 30 previous negative specimens were tested greater than 72 hours after testing with the CDC assay. In addition, a combination of 15 sputum/tracheal aspirate specimens (13 positive and 2 negative) were tested on the DIA, NDX, and GX instruments within 24 hours of each other. All NP/nasal swab and sputum/tracheal aspirate specimens were held at 2-8 °C until testing was completed.

### Platform Methods

#### CDC

Total RNA from 200 microliters (ul) of patient specimen was extracted with the NucliSENS ezMAG platform (BioMerieux, Durham, NC) according to manufacturer’s protocol with a final elution volume of 50 ul. For each specimen, three separate master mixes including primer and probe sets to detect the N1, N2, and RNaseP targets (Integrated DNA Technologies (IDT), Coralville, Iowa) using the TaqPath™ 1-Step RT-qPCR Master Mix, CG (ThermoFisher, Waltham, MA) were prepared with a final reaction volume of 15 μL of each master mix and 5 μL of extracted RNA. All reactions were performed in 96-well plates on a Roche LC 480 or z480 light cycler (Roche, Basel, Switzerland) according to the CDC procedure including a previously negative patient specimen (extraction and RNaseP internal controls), fresh UTM and a SARS-CoV-2 N1, N2, and RNaseP RNA positive controls (IDT). Patient specimens were considered positive if both N1 and N2 targets were detected.

#### Diasorin

Fifty ul of patient specimen and 50 ul of mastermix provided by the manufacturer was added to each well of the Simplexa COVID-19 Direct disc real time polymerase chain reaction assay (rtPCR) and performed according to manufacturer’s protocol on the Diasorin Liaison instrument (Diasorin, Saluggia, Italy). Each reaction well of the disc included an RNA internal control to determine PCR failure and/or inhibition. Results were analyzed using the LIAISON^®^ MDX Studio software supplied by the manufacturer and were considered positive if either one of the S or ORF1ab genes were detected.

#### NeuMoDX

Five-hundred microliters of UTM was added to 500 ul of lysis buffer provided by the manufacture (NeuMoDx, Ann Arbor, MI) and placed onto the automated rtPCR extraction/detection instrument according to manufacturer’s protocol. Results were analyzed using on-board instrument software and patients were considered positive if either the N or Nsp2 genes were detected.

#### Cepheid

Five-hundred microliters of UTM was added to the rtPCR test cartridge and placed onto the platform according to manufacturer’s protocol. GenXpert platform (Cepheid, Sunnyvale, CA, USA). Results were considered positive if either the E or N2 genes were detected. Sputum/Tracheal aspirates were diluted 1:4 in saline prior to testing and processed as previously described above for NP/nasal swabs in UTM on the DIA, NDX, and GX instruments only.

### Data analysis

Since the CDC/IDT assay was the first assay to obtain EUA approval it was considered the “gold standard” for the analysis of results from the NP/nasal swabs. Threshold cT values for both targets from each platform were combined into a single data set for each platform therefore each platform could have a total of 40 possible targets detected for the positive specimens. All analyses of cT values from DIA, NDX, and GS were done in comparison to cT values from the CDC assay including regression analysis, Pearson’s correlation coefficient (Pearsons r), and Chi-square were performed using Microsoft Excel (Microsoft, Redmond, WA). For the sputum/tracheal aspirate specimens, similar data analysis was done; however, since these were not tested with the CDC assay, the DIA assay was considered the “gold standard”.

### Results

Qualitatively (Positive/Negative), nineteen of the 20 positive and all 30 negative specimens using the CDC assay matched using the DIA, NDX and GX assays for an overall test interpretation accuracy of 98.0%. For the NP/Nasal specimens, based on total 40 possible targets per platform, one of the two targets for both the DIA and GX platforms was missed for a single CDC positive specimen resulting in 2 false negatives targets, therefore target accuracy of the three instruments were 97.5%, 100% and 97.5. %, in DIA, NDX and GX assays, respectively. However, taking into consider that only one of the two targets needed to be detected to consider the results positive, there was 100% accuracy across all 4 platforms (data not shown). Regression analysis of threshold cT values demonstrated strong correlation between all three platforms compared to the CDC assay with R^2^ values for the DIA, NDX, and GX cT results of 0.978, 0.9774, 0.8555, respectively (Figure 1). Further analysis using Pearson’s r and Chi^2^ testing demonstrated a nearly linear relationship and no significant differences (Table 1). The average threshold cTs for all targets from the DIA and NDX platforms were 3.7 and 2.7 lower, respectively compared to the CDC assay and the GX cTs were approximately the same (0.3) (Figure 2). All 15 of the sputum/tracheal aspirate specimens were repeated on both the NDX and GX instruments for 100% agreement between the three platforms. Regression analysis showed strong correlation between the DIA and NDX with an R^2^ of 0.953; however, this was not quite as robust for the GX, R^2^ = 0.758 (Figure 3). The linear relationship between the three platforms using Pearson’s r and Chi2 testing was not significantly different (Table 1).

**Figure 1.**
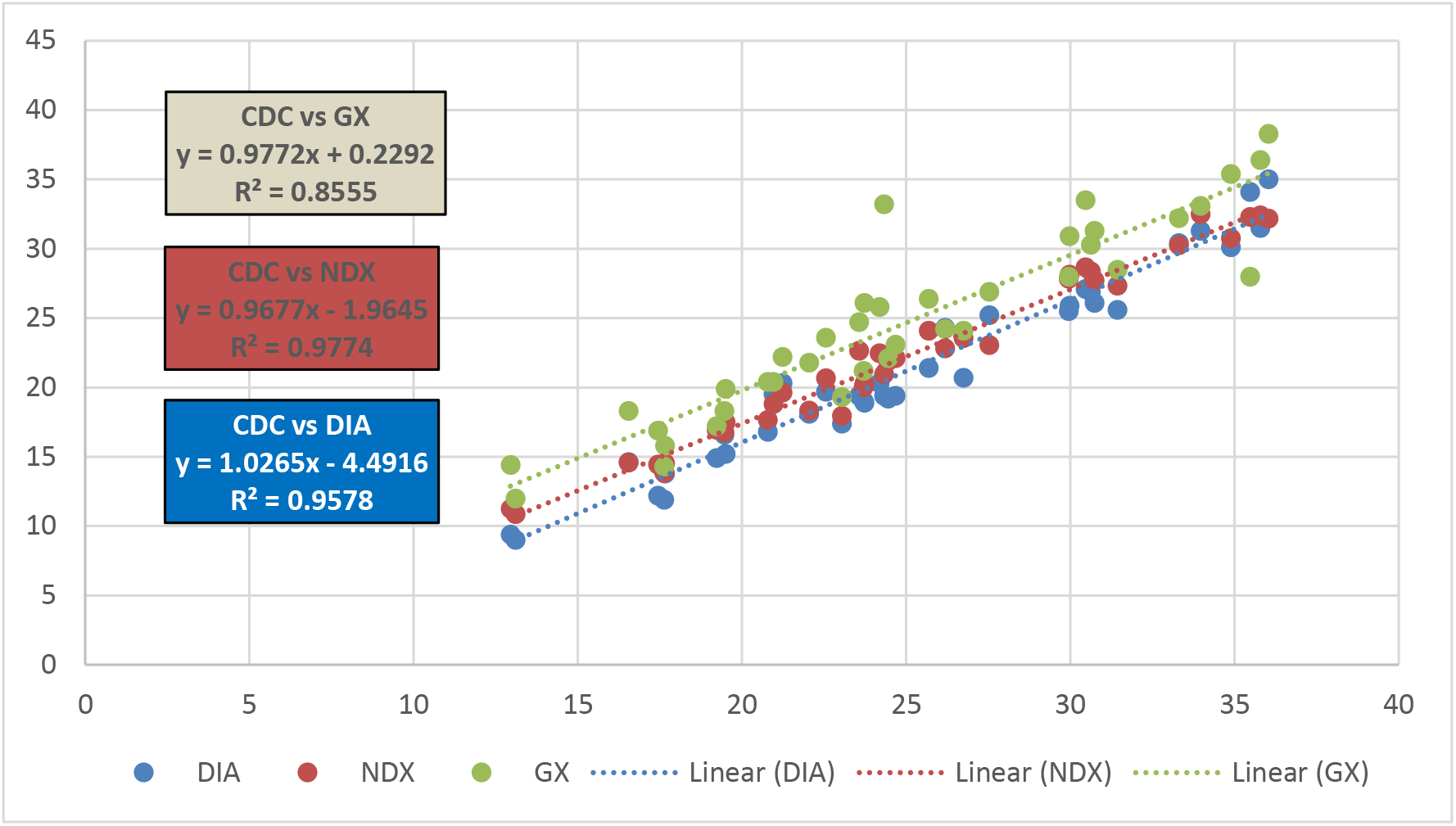
Comparison of threshold cTs obtained on the DIA, NDX, and GX platforms versus those from the CDC assay from NP/nasal swabs. The corresponding R^2^ values indicate strong correlation between the four instruments.

**Table 1.** Statistical analysis of the mean threshold cT from the DIA, NDX, and GX platforms compared the mean threshold cT from the CDC assay and the mean threshold cT from the NDX and GX compared to the DIA assay.

| NP/nasal swabs | cT Average | Pearson's r | Chi-Square |
| --- | --- | --- | --- |
| CDC Assay | 25.04 | NA | NA |
| DIA | 21.21 | 0.978 | 0.888 |
| NDX | 22.2 | 0.988 | 0.999 |
| GX | 24.6 | 0.924 | 0.999 |
| Sputum/Tracheal | cT Average | Pearson's r | Chi-Square |
| DIA | 22.9 | NA | NA |
| NDX | 23.5 | 0.976 | 1.00 |
| GX | 27.1 | 0.870 | 0.106 |

**Figure 2.**
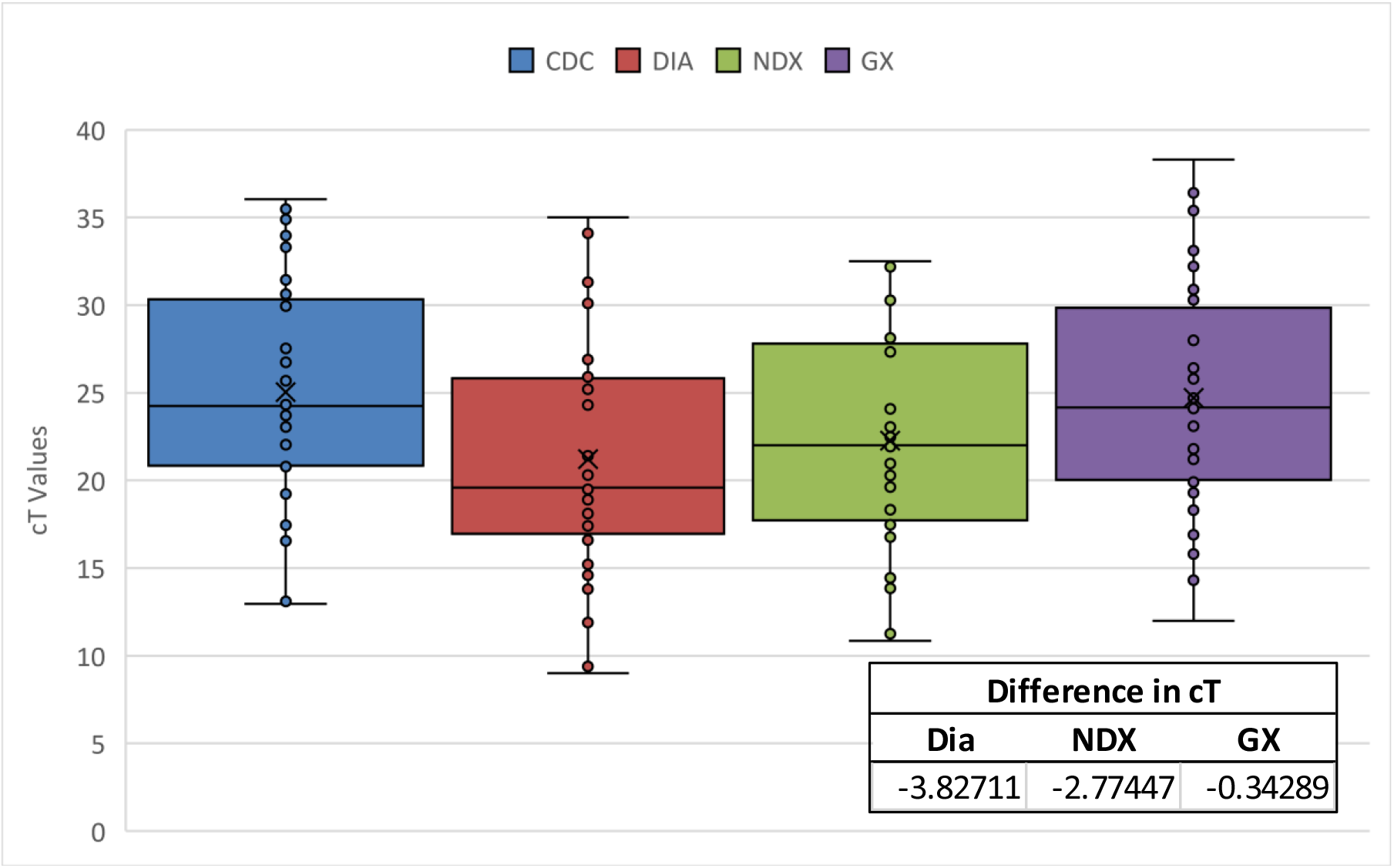
Overall threshold cT distribution of the means of all targets from each instrument obtained from NP/nasal swabs. DIA showed the lowest overall cT average of 21.2 followed by NDX (22.2), GX (24.6) and CDC (25.0). The inset shows that the DIA, NDX, and GX assays signaled positive 3.8, 2.77, and 0.4 cTs earlier than the CDC assay.

**Figure 3.**
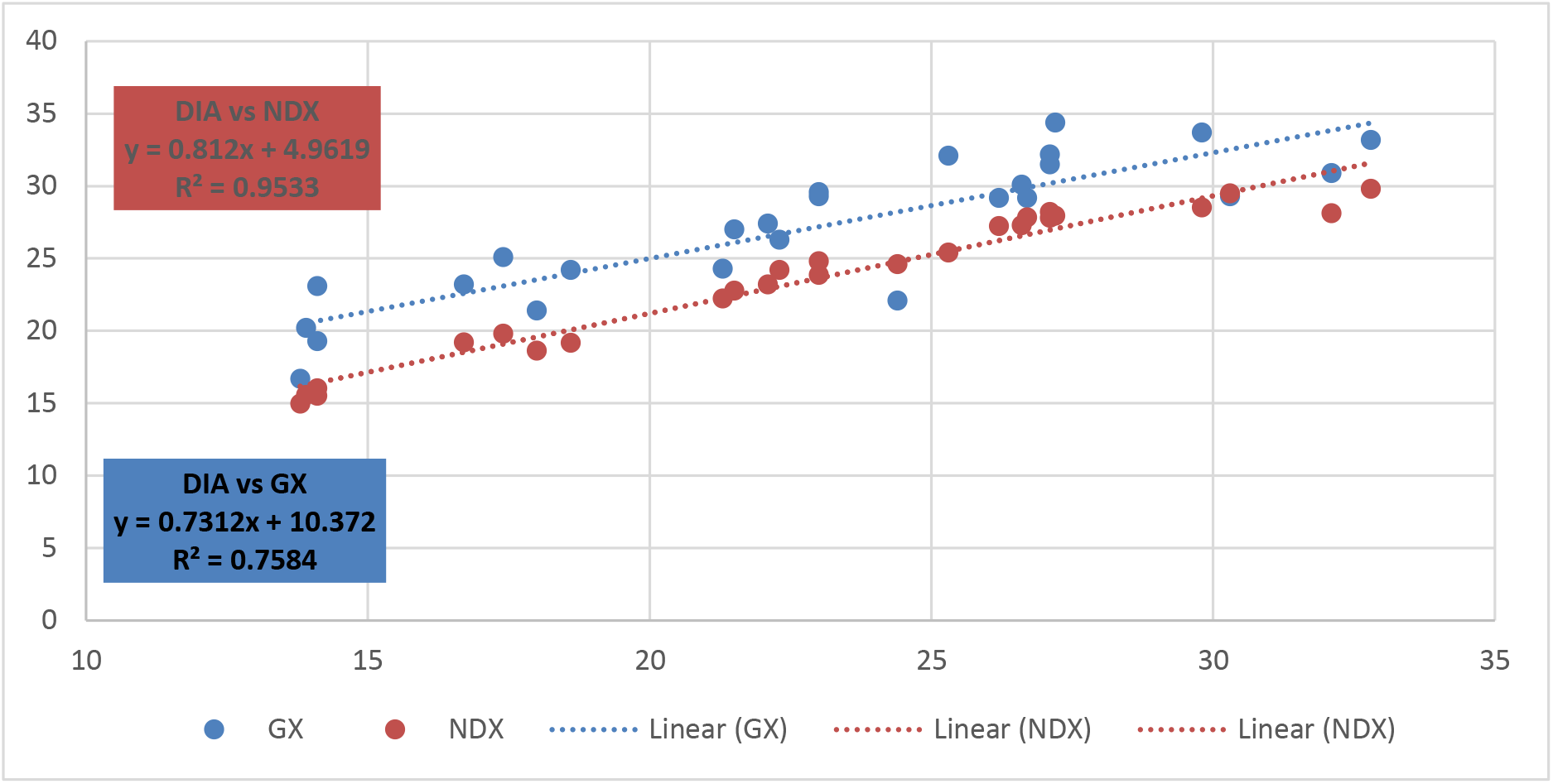
Comparison of threshold cTs obtained on the NDX and GX platforms compared to the DIA platform obtained from sputum/tracheal aspirate specimens. The corresponding R^2^ value for the NDX indicates a strong correlation compared to the DIA; however, not as strong for the GX instrument.

**Figure 4.**
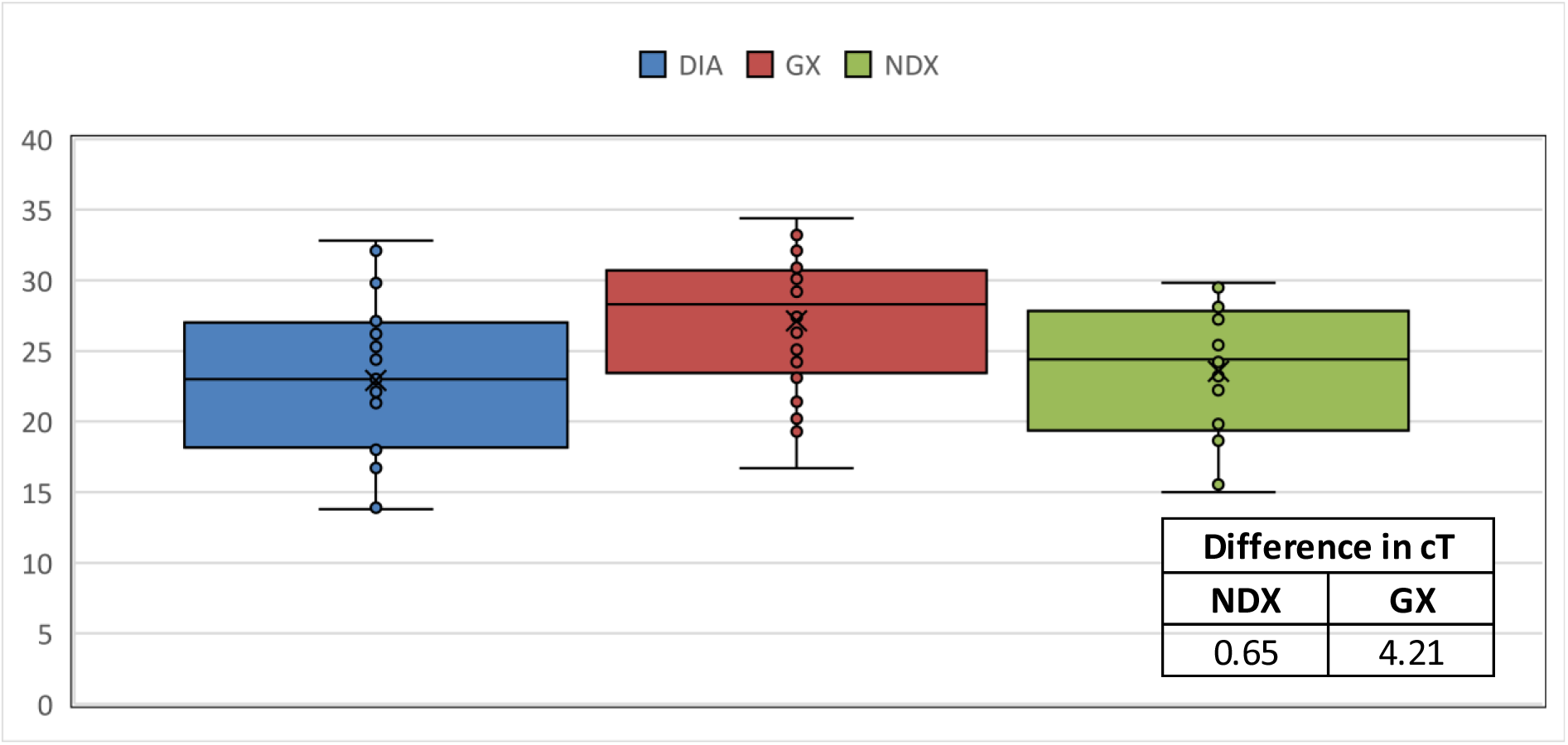
Overall threshold cT mean distribution of all targets from each instrument obtained from sputum/tracheal aspirate specimens. The DIA and NDX instruments were comparable at 22.9 and 23.5, respectively while the GX average was 27.4. The inset shows that the NDX and GX assays signaled positive 0.65 and 4.21 cTs later than the DIA assay.

### Discussion

The onset of the COVID-19, officially a pandemic by January 2019, made molecular diagnostic testing for SARS-CoV-2 an international race to develop solutions to testing and associated supply chains. In the United States, the FDA gave EUA approval to the CDC test platform in February and quickly began to approve EUA status to additional vendor tests and hospital laboratories to broaden testing availability. However, the markedly increased world-wide demand soon outstripped the supply leading to tense competition to maintain testing. Further, clinicians soon experienced unexpected negative PCR test results in clinical scenarios of active disease– a phenomenon that is well described in the literature (4). This led to questions of false positive PCR test performance versus sampling adequacy and reliability in relation to viral disease kinetics. We undertook this study to address potential differences in performance between the 4 SARS-CoV 2 rtPCR method implemented at Henry Ford Hospital. Here we aimed to compare the qualitative accuracy (positive/negative) and the relative analytic sensitivity of three of our new SARS-CoV-2 instruments to the CDC assay on a 4^th^ platform by looking at the overall threshold cT values of each. Qualitatively, the overall accuracy of the NP/nasal swabs using the 4 platforms was 98.0%, based on the assumption that missing one of the two targets is considered a negative result. This is similar to results reported by Rhoads et al in their comparison of the Diasorin and CDC assays (3). However, the EUA package insert for all 4 platforms indicates that only one of the two targets need be detected for either a positive or presumptive positive result. Therefore, based on this criterion our overall accuracy is 100% overall for NP/nasal swabs and sputum/tracheal aspirates. Since each assay has two targets, there were a total of 40 possible positive reaction targets in all specimens tested. However, for one specimen, a single target was missed on the DIA and GX platforms from one NP/nasal swab, therefore the accuracy was 95% for each of these methods. Of note, the DIA missed one of the targets as described above, despite having the strongest overall correlation to the CDC assay. However, the threshold cT for the CDC assay was near the end of the assay cycles (37.95 and 39.94/40) therefore, it was most likely a weak positive. As shown in Table 1, these platforms compared very well for both UTM and sputum/tracheal aspirates. Similarly, Zehn et al also demonstrated an overall strong correlation between the Diasorin, GenMark and CDC assays (5). For their study they obtained a kappa score analysis of 0.96, while our correlations between the platforms ranged from 0.92 – 0.97 and 0.88 – 0.99 using Pearson’s r and chi-square analysis respectively. Similar results were obtained using sputum/tracheal aspirates for the NDX (0.87) and GX (0.87) for the Pearson’s r analysis but did see a much lower value for the GX (0.1) versus the NDX (1.0) using chi-square; however, the difference was not significant. In addition, comparing threshold cTs using regression analysis resulted in R^2^ values from 0.85 – 0.97 for NP/Nasal swabs in UTM (Figure 1); however, while the R^2^ for sputum/tracheal aspirates on the NDX was 0.95, the R^2^ was only 0.75 for the GX, thus matching the previous analysis (Figure 3).

While there was strong correlation between the 4 platforms, we also wanted to evaluate the average threshold cT for positive results on each platform. For the NP/nasal swabs the average threshold cT was lowest for the DIA followed by the NDX and GX (Figure 2). Similar results were seen for the sputum/tracheal aspirates with DIA being the lowest followed by the NDX and GX platforms. For the NP/nasal swabs these averages on the DIA and NDX crossed 3-4 cTs earlier than the CDC assay while the GX was about the same (Inset Figure 2). Conversely, for the sputum/tracheal aspirates, the average threshold cTs were about the same or up to 5 cTs higher for the NDX and GX respectively compared to the DIA. This is interesting because the NDX and GX both require 500 ul of specimen; however, the DIA only requires 50 ul. A limitation of this study is the lack of limit of detection data for the DIA, NDX, and GX platforms. However, for the initial validation of the EUA CDC assay we determined that the limit of detection using plasmid-based target was 1 copy/ul and between 20-40 copies/ul using SARS-CoV-2 RNA. Historically it has been shown that every cT change of 3.3 equates to a 1 log10 difference in quantity for molecular assays. Therefore, it appears that based on the average cT values, the DIA was the most sensitive of the 4 platforms.

The DIA and GX assays offer different workflow options in comparison to the NDX platform. The NDX has two platform options - the 288 and 96 which can process approximately 200 and 90 samples for SARS-COV-2 PCR in 8 hours. The testing in this study was performed on the NeumoDx 288. The DIA and the GX assays on the other hand have relatively lower throughput but more rapid time to result (75 minutes and 55 minutes respectively). The DIA is capable of processing 8 samples/run/instrument while the GX throughput is dependent on the number of cartridge slots available per instrument with each sample being loaded individually. The NDX is typically batch loaded but does have a random-access priority testing option.

Due to the emergent need for highly sensitive methods to detect SARS-CoV-2, it has become standard practice in many clinical microbiology laboratories to employ multiple platforms to accommodate a high volume of testing. The obvious question that one could ask is which is the “best” assay to use? This analysis has demonstrated that despite use of different targets, all 4 assays were able to detect SARS-CoV-2 from multiple specimen types with relatively high accuracy and threshold cT correlation. We are confident that this rtPCR testing reproducibility demonstrated here validates the clinical use of numerous test platforms for detection of SARS-CoV-2.

